# Salivary *AsHPX12* influence pre-blood meal associated behavioral properties in the mosquito *Anopheles stephensi*

**DOI:** 10.1101/2020.06.12.147959

**Authors:** Seena Kumari, Tanwee Das De, Charu Chauhan, Jyoti Rani, Sanjay Tevatiya, Punita Sharma, Kailash C Pandey, Veena Pande, Rajnikant Dixit

## Abstract

In the adult female mosquito, successful blood meal acquisition is accomplished by salivary glands, which releases a cocktail of proteins to counteract vertebrate host’s immune-homeostasis. However, the biological relevance of many salivary proteins remains unknown. Here, we characterize a salivary specific Heme peroxidase family member HPX12, originally identified from *Plasmodium vivax* infected salivary RNAseq data of the mosquito *Anopheles stephensi*. We demonstrate that dsRNA silencing mediated mRNA depletion of salivary *AsHPX12* (80-90%), causes enhanced host attraction but reduced blood-meal acquisition abilities, by increasing probing propensity (31%), as well as probing time (100–200s, *P<0.0001*) as compared to control (35-90s) mosquitoes group. Altered expression of the salivary secretory and antennal proteins may account for an unusual fast release of salivary cocktail proteins, but the slowing acquisition of blood meal, possibly due to salivary homeostasis disruption of *AsHPX12* silenced mosquitoes. A parallel transcriptional modulation in response to blood feeding and *P. vivax* infection, further establish a possible functional correlation of *AsHPX12* role in salivary immune-physiology and *Plasmodium* sporozoites survival/transmission. We propose that salivary HPX12 may have a vital role in the management of ‘pre- and post’-blood meal associated physiological-homeostasis and parasite transmission.

**Graphical abstract:** Figure 1:
Schematic representation of mosquito’s blood meal acquisition and upshot on blood-feeding after silencing of salivary gland HPX-12. **(A)** After landing over host skin, mosquito mouthparts (proboscis) actively engaged to search, probe, and pierce the skin followed by a rapid release of the pre-synthesized salivary cocktail, which counteracts the host homeostasis, inflammation, and immune responses, during blood meal uptake. **(B)** Silencing of HPX-12 disrupts salivary gland homeostasis, enhancing mosquito attraction, possibly by up-regulating odorant-binding proteins genes-OBP-7,10 and OBP-20 expression in the Olfactory System. However, HPX-12 disruption may also cause significant effects on pre-blood meal associated probing abilities, which may be due to fast down-regulation of salivary cocktail proteins such as Anopheline, Apyrase, D7L proteins.

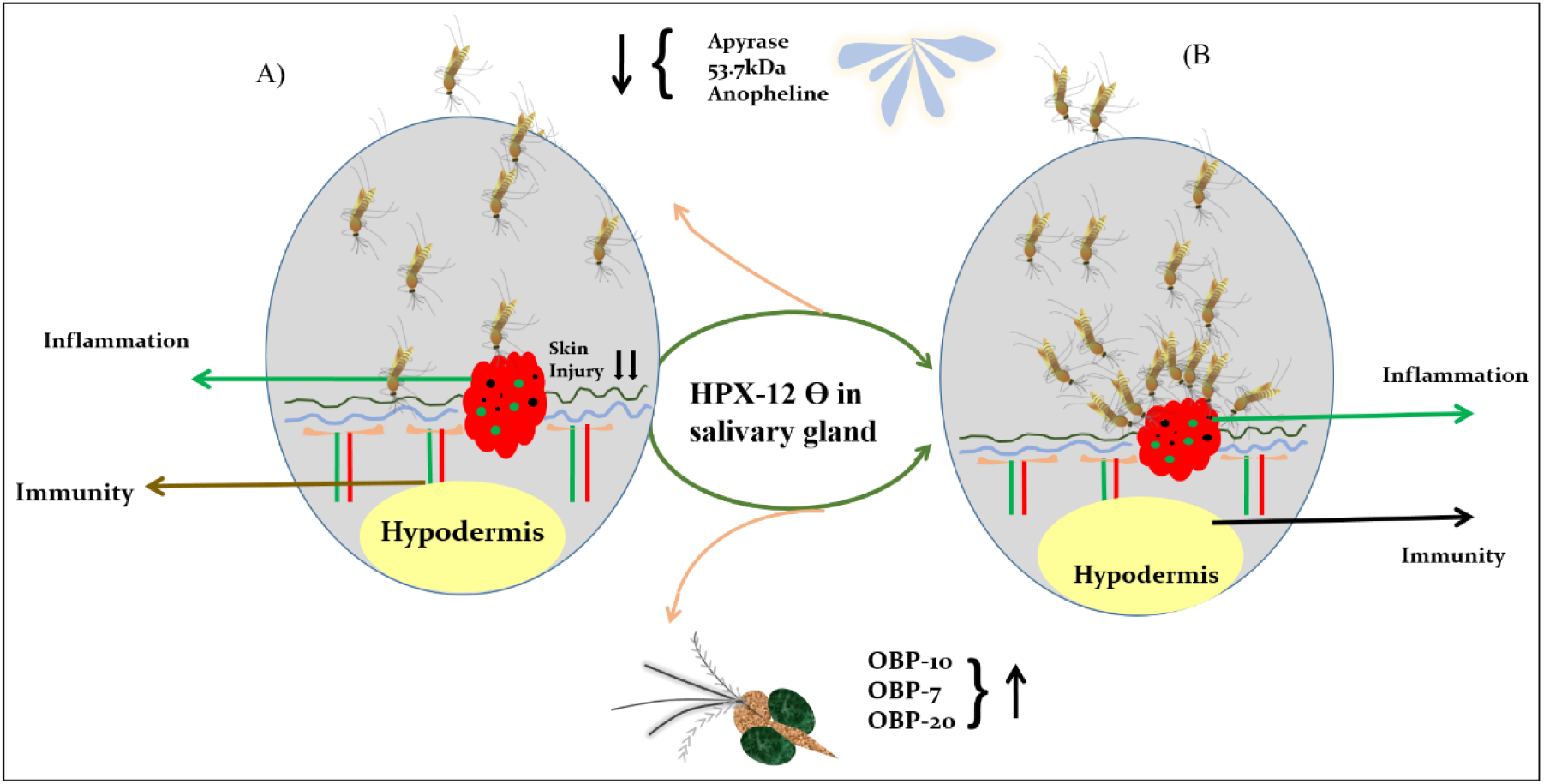

## 1. Introduction

Adult mosquitoes of both sexes relies on nectar sugar for their regular metabolic energy production (1–4). However, an evolutionary adaptation of host-seeking and blood-feeding behavior in the adult female mosquitoes are imperative for their reproductive success. If taken infectious blood meal, it may also affect their ability to transmit several infectious diseases (5–10). The harmonious actions of the neuro-olfactory system drive mosquito’s successful navigation towards a vertebrate host (11), but it is the salivary gland that facilitates rapid blood meal uptake from the respective host (12). The mosquito’s salivary cocktail contains crucial bioactive molecules having anti-homeostatic (13–15), anti-inflammatory, and immunomodulatory properties (16–19), which counteract the host defense for rapid blood meal acquisition, usually in less than two minutes.

For the past two decades, several salivary glands encoded factors have been identified from different mosquito species (20–23). But the nature and function of a salivary cocktail while changing active physiological status from sugar-to-blood feeding remains largely unknown. Substantial evidence shows that salivary gland secretory proteins are rapidly depleted upon blood-feeding (23–25). Our recent study also demonstrates that mosquito salivary glands gene expression switching is key to manage meal specific responses. Additionally, we also observed that the first blood meal not only modulates the molecular and cellular responses but also causes persistent changes in the salivary gland morphology(26). Though we correlate the pre and post blood meal associated changes are pivotal to maintain salivary gland homeostasis, but the regulatory mechanism remains unexplored.

Heme containing peroxidase enzymes, having conserved function throughout vertebrates and invertebrates taxa, play a crucial role in the maintenance of cellular and physiological homeostasis by scavenging free radicals (NOS/ROS)(27–30). Out of a total of 889 putative insect heme peroxidases indexed in the NCBI database, at least 39 heme peroxidases have been predicted from blood-feeding mosquitoes, which are distributed to six highly conserved HPX lineages. Though salivary peroxidase constitutes important bioactive vasodilator molecules, their functional role has not been established (31).

Here, we demonstrate that *AsHPX12*, a heme peroxidase homolog, abundantly expresses in the salivary glands of *Anopheles stephensi* mosquito. Using dsRNA mediated gene silencing experiments, we show that salivary *AsHPX12* dysregulation significantly impairs pre blood meal associated behavioral properties such as delay in probing time and blood meal acquisition propensity of the mosquitoes. Possibly, this is caused by salivary physiological homeostasis disruption and altered expression of olfactory and salivary proteins of the *AsHPX12* mRNA depleted mosquitoes. Furthermore, a significant transcriptional modulation in response to blood feeding and *P. vivax* infection, suggests salivary *AsHPX12* may have a unique role in the maintenance of salivary gland physiology, and *Plasmodium* sporozoite survival and transmission.

## 2. Material and Methods

### 2.1 Mosquito Rearing and Maintenance

Indian strain of *An. stephensi* mosquito was reared and maintained in the insectary at the temperature of 28 ± 2 °C, the relative humidity of ∼80% with 12h light-dark cycle as mentioned previously (32)(33). The adult mosquitoes were fed daily on sterile sugar solution (10%) using a cotton swab throughout the experiment. All protocols for rearing and maintenance of the mosquito culture were approved by the Institute Animal Ethics Committee (ECR/NIMR/EC/2012/41).

### 2.2 Sample collection and RNA extraction

Experimentally required developmental stages or tissues were dissected and pooled from the cold anesthetized adult female mosquitoes under different physiological conditions. To examine the tissue-specific expression of target genes, selected tissues such as hemocyte, spermatheca, olfactory system were dissected from 3-4-day old naïve sugar-fed mosquitoes. Salivary glands were dissected from 25 blood-fed (Rabbit) mosquitoes and collected at selected time points (3hr, 24hr, 48hr, 72hr, 10 days, and 14 days). For the collection of salivary glands infected with *P. vivax* sporozoites, 3-4D old *An. stephensi* mosquitoes were fed on the *P. vivax* infected patient’s blood (∼2% gametocytemia) through pre-optimized artificial membrane feeding assay (34). The confirmation of the *P. vivax* infection was done by staining the midgut with 5% mercurochrome to visualize the oocysts after 4 days post-infection (DPI), as described earlier (34). After confirmation, ∼20-25 mosquitos’ Salivary gland dissected at 9-12DPI and 12-14DPI, respectively. Total RNA from the salivary gland, midgut, and other tissues were isolated using a standard Trizol method as described previously (26).

### 2.3 cDNA preparation and gene expression analysis

Isolated ∼1µg total RNA was utilized for the synthesis of the first-strand cDNA using a mixture of oligo-dT and random hexamer primers and Superscript II reverse transcriptase as per the described protocol (Verso cDNA synthesis Kit, Cat#AB-1453/A, EU, Lithuania; (26). For differential gene expression analysis, routine RT-PCR and agarose gel electrophoresis protocols were used. The relative abundance was assessed by SYBR green qPCR master mix (Thermo Scientific), using Illumina Eco-Real Time or Bio-Rad CFX96 PCR machine. PCR cycle parameters involved an initial denaturation at 95°C for 15 min, 40 cycles of 10 s at 95°C, 15 s at 52°C, and 22s at 72°C. After the final extension steps melting curves were derived. Each experiment was performed in three independent biological replicates. The relative quantification results were normalized with an internal control (Actin), analyzed by 2–^ΔΔ^Ct method, and statistical analysis was performed using Origin 8.1.

### 2.4 dsRNA mediated gene silencing assays

For the knocking down of *AsHPX12*, dsRNA primers carrying T7 overhang were synthesized as listed in the supplemental table (ST2). The amplified PCR product was examined by agarose gel electrophoresis, purified (Thermo Scientific Gene JET PCR Purification Kit #K0701), quantified, and subjected to double-stranded RNA synthesis using Transcript Aid T7 high-yield transcription kit (Cat# K044, Ambion, USA), and bacterial dsrLacZ gene was used as control. About ∼69 nl (3 µg/ul) of purified *dsRNA* product was injected into the thorax of cold anesthetized 1–2-day old female mosquito using nano-injector (Drummond Scientific, CA, USA). The silencing of the respective gene was confirmed by quantitative RT-PCR after 3-4 -days of dsRNA injection.

### 2.5 Blood-feeding assays

To track the possible role of *AsHPX12* in blood-feeding both control (*Lac*Z injected) and test (*AsHPX12* injected) mosquitoes, were offered rabbit blood meal after 4 days of the dsRNA injection. The mosquitoes were allowed to feed for 20 min on rabbit (shaved pinna), and thereafter we scored the number of mosquitoes that had fed. For the statistical analysis of blood-feeding propensity (percentage of mosquitoes that probed within a fixed time-period), the feeding phenotype of silenced mosquitoes was compared with the respective control group (35,36). For probing assay also, the control and *AsHPX12* silenced mosquitoes were starved for 4-5 hours before exposure to vertebrate rabbit host (saved pinna), and calculated probing time (the initial insertion of the proboscis into the skin to the initial engorgement of blood) as described by Lombardo *et al*., (37). Briefly, LacZ and HPX-12 silenced mosquitoes (30 in each batch) were released into the modified Olfactometer which comprised two arms, each ending in two independent tests and control chamber (length 44 cm, width 36 cm). Before the initialization of the experiment, both groups of mosquitoes were allowed to acclimatize for 30 minutes within the chamber. After acclimatization two rabbits were kept into the arms of the Olfactometer and probing time was calculated.

### 2.6 Trypan blue staining

After participating in experiments, females were cold anesthetized, and placed in a drop of phosphate-buffered saline (PBS) over microscopic glass slide, and examined under dissecting microscope (16×). Using micro-needles salivary gland tissue was gently removed by pulling out from thorax. Individually dissected salivary glands from control or test mosquito group were incubated in trypan blue 0.4% dye (1:1) for 1-2 minutes at room temperature. Salivary glands were washed three times with PBS and transferred to a new glass slide in a fresh drop of PBS. The subsequent observations were made under a compound microscope (100×), to visualize the ultrastructure of stained salivary glands. The images of tissue were capture with the assistance of Smart-Phone Camera (with zoom option/12X), stabilized on the eyepiece of the microscope. The final images of both control & test groups were captured in identical magnifying parameters and further processed through professional software (Microsoft Photo Edit, version 2020.19111.24110.0).

### 2.6 *In-silico* Bioinformatics analysis

Phylogenetic trees were prepared with selected peroxidase protein sequences by maximum likelihood (ML) method in MEGA X program as described previously (38). We aligned all our selected peroxidase protein sequences by ClustalW algorithm where the reliability of the branching was tested by 1000 bootstrap value of the replicates. The processed Phylogenetic tree was examined based on clusters and nodes formed.

### 2.7 Statistical analysis

Statistical analysis was performed using Origin8.1. All these data were expressed as mean ± SD Differences between test samples and their respective controls were evaluated by paired Student’s *t*-test and one-way ANOVA test considered to be significant if the *p*-value was less than 0.05. Each experiment was performed at least thrice to validate the findings.

## 3. Results

### 3.1. Identification, annotation, and Phylogenomics Analysis of *AsHPX-12*

Our recent RNAseq study demonstrates that *P. vivax* sporozoites significantly modulates the molecular architecture of the mosquitoes’ salivary glands (39). To identify the transcripts having a crucial impact on salivary physiology and antioxidant defense responses, we selectively cataloged at least six transcripts encoding heme-peroxidase homologous proteins (Table-1) from transcriptomic data (38). Relatively a high read count of HPX12 than other members of the heme peroxidase family prompted us to investigate its possible role in the mosquito salivary physiology. The BLASTX analysis of the selected 1806bp long HPX12 transcript showed 99% identity with the HPX12 homolog of *Anopheles gambiae* (AGAP029195), having conserved Animal heme peroxidase domain (Fig.2a). A homology search against *An. stephensi* database further predicts full-length *AsHPX12* (ASTE015997) is a 1905 bp long single-copy gene. It encodes a protein of 595 amino acid (aa) having 7 motifs including Amidation, Casein kinase-II phosphorylation, protein kinase C and N-myristoylation site, cAMP, and cGMP dependent protein kinase phosphorylation site.

**Table1.** Number of transcripts encoding distinct Heme peroxidases retrieved from the RNAseq database of blood-fed (control) and *Plasmodium vivax* infected salivary glands of *An. stephensi*

| S.No. | Assembled contigs | Length | Best match to AG_immunoPDB | E-values | Read Count (control) | Read Count (infection) |
| --- | --- | --- | --- | --- | --- | --- |
| 1. | CDS_4640_Transcript_26174 | 1821 | HPX-12 | 0.00 | 17895 | 62566 |
| 2. | CDS_7327_Transcript_59398 | 591 | HPX-10 | 3E-84 | 10 | 30 |
| 3. | CDS_6192_Transcript_40923 | 369 | HPX-4 | 3.00E-13 | 9 | 14 |
| 4. | CDS_5032_Transcript_2704 | 450 | Duox | 2.00E-10 | 374 | 969 |
| 5. | CDS_4329_Transcript_25430 | 309 | HPX-2 | 2E-32 | 6 | 24 |
| 6. | CDS_6665_Transcript_48378 | 300 | HPX-3 | 5E-29 | 7 | 44 |

Multiple sequence alignment of putative HPX members from insects and mosquitoes indicated a high degree of sequence conservation within the *Anopheles* species, where *AsHPX12* showed the highest identity (85%) match to *An. gambiae* (Fig.2b, ST-1). The phylogenetic analysis resulted in the formation of two independent clades, where each clade defines separate lineage for animal heme peroxidase, which is further divided into specific subclades such as HPX2, HPX4, HPX6, HPX8 and Duox (Fig.2c). Additionally, selective Phylogenetic analysis of *AsHPX12* homolog members among vector and non-vector mosquito species (ST1) revealed *AsHPX12* forms an independent clade with *An. funestus* (Fig.2d).

**Figure 2:**
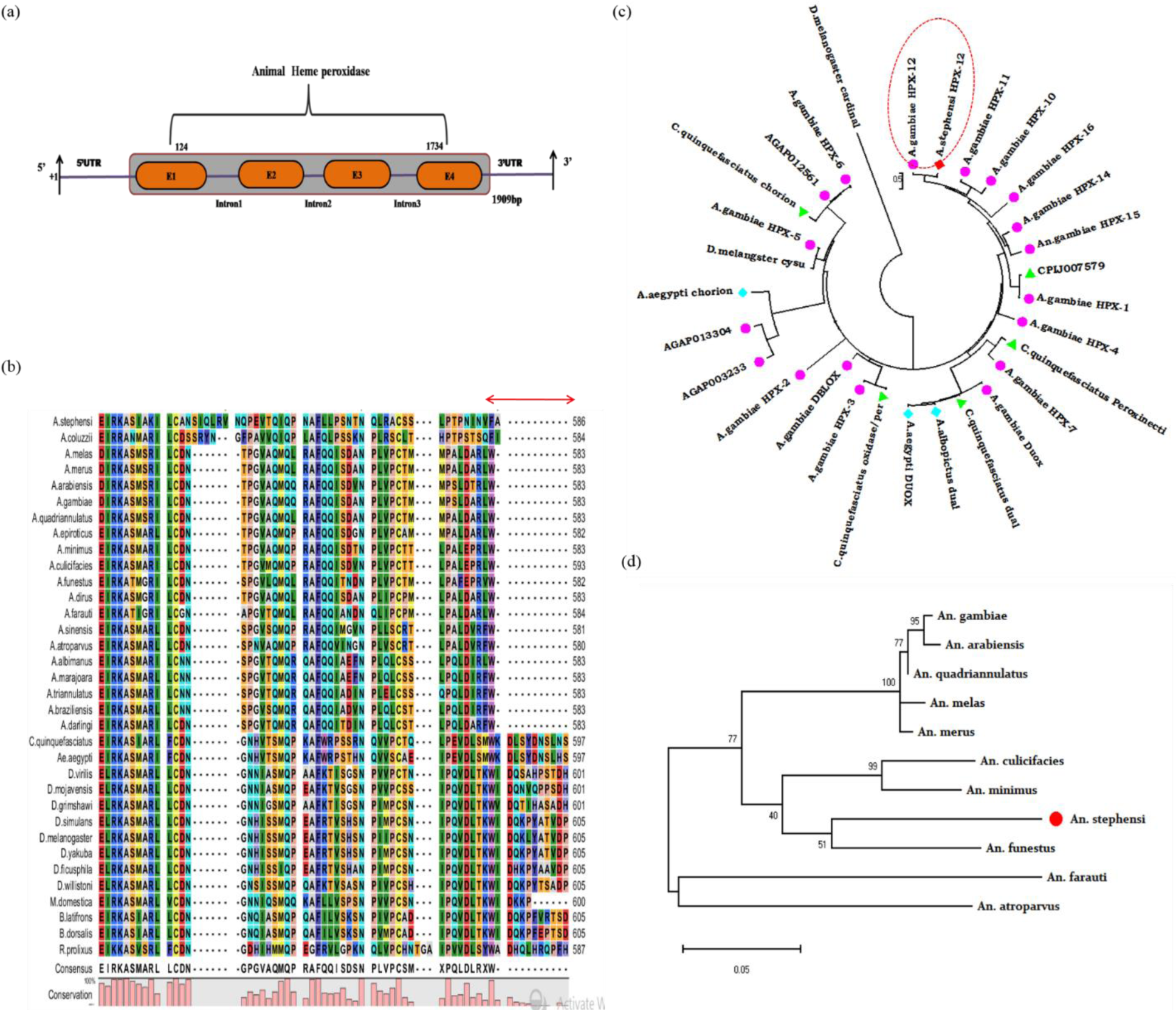
Genomic organization and molecular characterization of *An. stephensi HPX*-12: **(a)** Schematic representation of the genomic architecture of *AsHPX12*. Four brown color boxes (E1-E4) represent exons and +1 indicates translation initiation site. A 50 bp UTR region is present on both 5’ and 3’ end of the transcript; **(b)** Multiple sequence alignment showed *Anopheles* species lack a stretch of 11 AA at the 3’end, which is present in other insects except mosquito i.e. Drosophila; **(c)** Phylogenetic relationship of HPX family proteins within the insect’s community. The pink circle represents *Anopheles* species, light blue color circle denote *Aedes* mosquito species and green color triangle shows *Culex* species**;(d)** Phylogenetic relationship of our selected hpx-12 with other insects showed its association with *An. funestus.*

### 3.2 *AsHPX12* abundantly expresses in the salivary glands of adult mosquitoes

To predict the possible role of the identified salivary *AsHPX12*, first, we performed developmental expression analysis. Our initial RT-PCR based data analysis revealed *AsHPX12* ubiquitously expresses throughout the developmental stages of the mosquitoes (Fig.3a). A comparative tissue-specific transcriptional profiling by real-time PCR revealed that HPX12 abundantly expresses in the salivary gland than other tissues of in 3-4 day old naïve mosquitoes. Taken together we hypothesize that *AsHPX12* may have a salivary specific role in the regulation of mosquito’s blood-feeding associated behavioral properties.

**Figure: 3.**
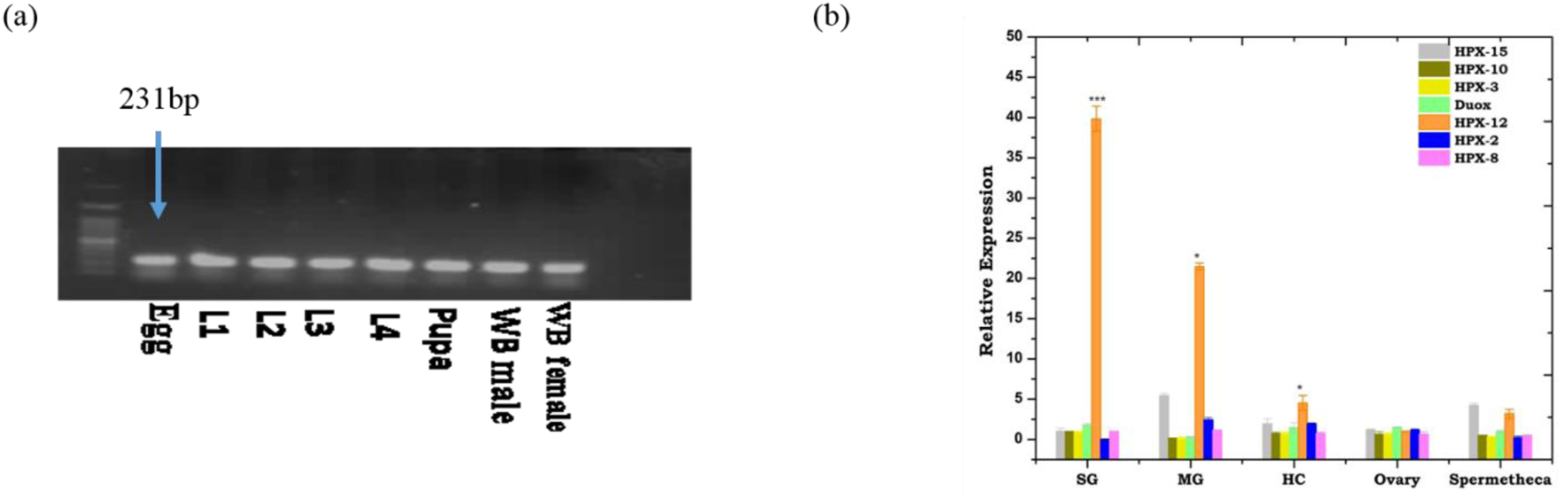
(a) RT-PCR based developmental expression analysis of *AsHPX12* in *An. stephensi* mosquito aquatic stages L1(larval stage1), L2(larval stage2), L3(larval stage3), L4 (larval stage 4) and whole body in male and female (n=10; N=3); (b)Tissues specific expression kinetics of HPX family in female mosquito SG: Salivary glands; MG: Midgut; HC: Hemocytes, OV: Ovary; SP: Spermetheca (n=25; N=3). Statistical significance was analysed using student’s t-test, (*\*P < 0.05; **P < 0.001; ***P < 0.0001*). (*n*=represents number of mosquito pooled for sample collection; *N*= number of replicates)

### *AsHPX-12* regulates pre-blood meal associated behavioral responses

To test the above hypothesis, we first examined the knockdown effect of *AsHPX12* on the mosquito’s pre-blood meal associated behavioral properties influencing the blood meal acquisition process. While performing behavioral assays, unexpectedly, we observed an increased host attraction of *AsHPX12* knockdown mosquitoes towards vertebrate host (ST3). However, interestingly, we also observed that the silencing of HPX12 (Fig. 4a, *p*<0.002)), significantly increases probing propensity by 31% (*p*<0.0067) and probing time to (∼100–200s, *P*<0.0001) as compared to ∼(35-90s) seconds of the control mosquitoes group (Fig. 4c; ST-3). Finally, when stained with trypan blue, which is a dye used to stain dead cells, the HPX12 silenced mosquito salivary glands showed an abnormal morphological disruption than control mosquito salivary glands (Fig. 4b.). Together, these data suggested that salivary HPX12 disruption may impair salivary physiology, resulting in altered functions during the blood meal acquisition process. It is well known that host-seeking behavioral property is regulated by the concerted actions of the odorant-binding proteins (OBPs) and odorant receptors (ORs). In a bid to check the correlation of salivary *AsHPX12* with pre-blood meal associated mosquitoes’ olfaction, we evaluated and compared the expression of selected OBPs/ORs in the naive and *HPX12* silenced mosquito’s salivary gland and olfactory tissues(4d). Among all the tested OBPs, exceptionally, we observed a significant up-regulation of OBPs, (*p*<0.01), but down-regulation of salivary apyrase (p<0.005), in both the salivary gland and olfactory tissues in *AsHPX12* silenced mosquitoes (Fig 4d,e). While the expression of odorant receptor remains unchanged in the olfactory as well as salivary glands (Fig.4d,e).

**Figure: 4.**
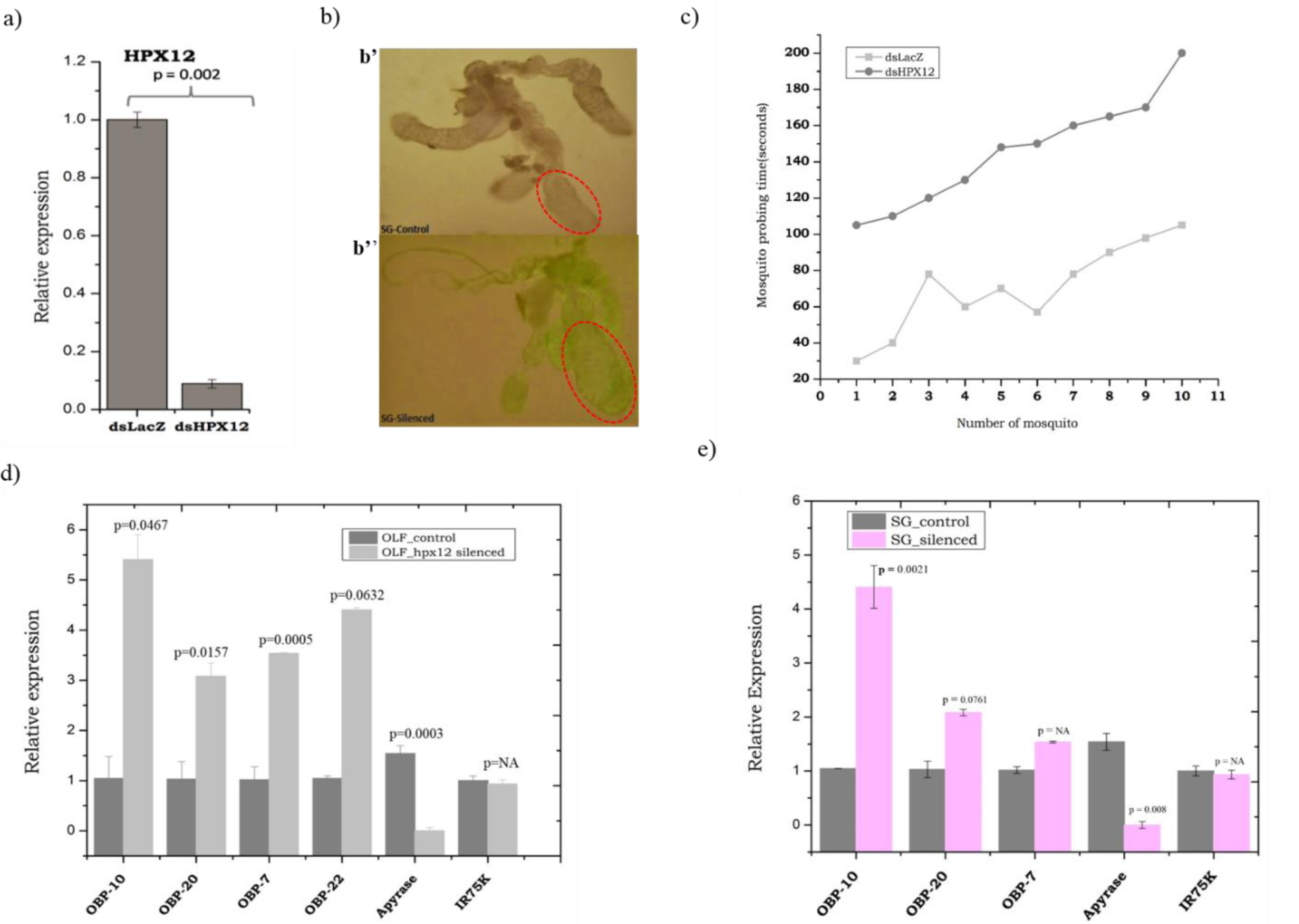
Alteration of molecular and behavioral properties in *AsHPX12* knockdown mosquitoes. **(a)** The relative abundance of *AsHPX12* mRNA in the salivary gland of control (dsLacZ injected) and silenced (dsHPX-12 injected) mosquitoes (p<0.002) (n=25, N=3); **(b)** Salivary glands of 5-day-old adult female control and HPX12 silenced mosquitoes were stained with trypan blue; ***(b’)*** a normal salivary gland from a 4 –day-old naive adult female mosquito; ***(b”)*** an abnormal salivary gland showing aberrant/distorted distal-lateral lobes of Hpx12 silenced mosquitoes; (marked red circle indicates the comparative morphological changes in the lobes). **(c)** Measurement and comparison of probing times of control and HPX12 silenced mosquitoes (N=3). Each dot corresponds to one female mosquito. The probing time is defined as the time taken from the initial insertion of the mouthpart into the skin until the initial observation of the ingestion of blood in the abdomen. Probing times were significantly longer in HPX12 silenced mosquitoes than in control mosquitoes. Data from three independent experiments using separate generations of mosquitoes were pooled. The number and ratio of blood-fed mosquitoes within 200 seconds, significance (*p*<0.000692), was calculated by one-way ANOVA. **(d)** comparative transcriptional profiling of the olfactory genes in the olfactory system (Consisting of Antenna, maxillary pulp and proboscis control vs HPX12 silenced mosquitoes (n=25, N=3). Transcript details are as follows: OBPs 10 (Odorant Binding Protein 10), OBP20, OBP7, OBP22, and OBP receptor IR75K (Ionotropic receptor 75K); Statistical significance was analyzed using student’s *t-*test (*\*P<0.05; **P<0.001; ***P<0.0001*). (*n*=represents number of mosquito pooled for sample collection; *N*= number of replicates).

### Blood meal and *P. vivax* sporozoite boost salivary *AsHPX-12* expression

Next, to evaluate how blood meal affects the expression of heme-peroxidase family proteins, we profiled and compared time-dependent transcriptional responses of all six HPX related family members in the salivary gland of blood-fed mosquitoes. Compared to other members, HPX12 showed a gradual induction within 6hrs of blood-feeding, which further increases to >16 fold (*p*<0.0025) by 24hrs after blood-feeding, and gradually cease to basal level after 72 hr of blood-feeding (Fig. 5a). Since, earlier studies demonstrate that a gut-specific *AsHPX15*, may favor *Plasmodium* development by the formation of a crosslinking mucin layer at the luminal side of the gut epithelium (40), we also tested whether *Plasmodium* infection also modulates *HPX12* responses in the mosquito salivary glands. Detailed transcriptional profiling showed a gradual elevation of *AsHPX*12 level in response to salivary invaded *P. vivax* sporozoite infection (Fig. 5b).

**Figure 5.**
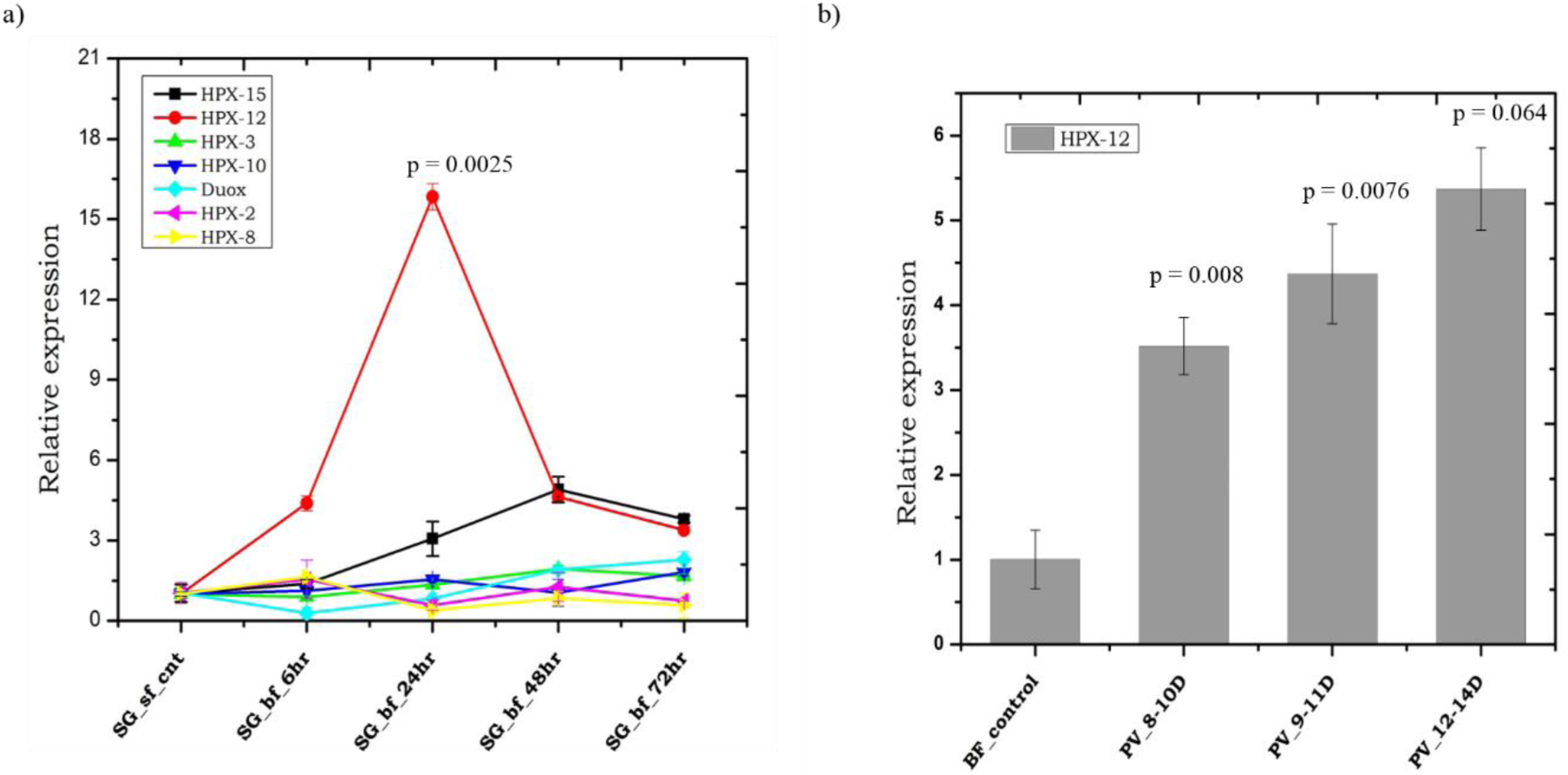
Transcription kinetics of HPX family members under different pathophysiological conditions of normal blood-meal time series and *Plasmodium vivax* infection. **(a)** Relative expression profiling of HPX family (hpx12,3,10,8 and hpx15, duox) members during blood meal time-series experiment; Salivary gland (SG) were collected from naive sugar-fed adult female mosquito and post blood-meal (6hr,24hr,48hr,72hr) (n=25, N=3); **(b)** Transcriptional profiling of Hpx12 in response to *P. vivax* infection, collected at selected time points (8-10D, 9-11D and 12-14D) days post-infection. Salivary Glands (SG) of age-matched normal blood-fed severed as control (n=25, N=3). Statistical significance was analysed using student’s *t-*test (*\*P<0.05; **P<0.001; ***P<0.0001*). (*n*=represents number of mosquito pooled for sample collection; *N*= number of replicates).

## Discussion

The evolution of conserved anti-oxidant defense system enzyme is necessary to maintain physiological homeostasis, but its role in salivary physiology of hematophagous insects remains unknown. Delayed probing and blood meal acquisition time can trigger awareness in the vertebrate hosts and may result in feeding challenge, or even death of the mosquito. Hence, an evolutionary adaptation of faster probing and feeding favored reducing vector-host contact duration and increased chances of blood feeder survival. Here, we identify and characterize a salivary specific *AsHPX12* transcript encoding heme-peroxidase enzyme, from malaria vector *An. stephensi*. We demonstrate *AsHPX12* is key to regulate the salivary homeostasis, and mRNA depletion by dsRNA silencing significantly impairs pre-blood meal associated host-seeking abilities of probing propensity and probing time, influencing blood meal acquisition process.

The navigation trajectory during active host-seeking is achieved by coordinated neuro-olfactory actions to find and locate a suitable vertebrate host (12), and once located the desired host, the salivary actions facilitate rapid blood meal acquisition process. Several studies suggest that during blood feeding the salivary cocktail composition is rapidly depleted (24), but how the host-odor activated salivary glands manages ‘prior and post’ blood-meal associated changing physiologies is largely unknown (26). We aimed to screen and test the possible role of HPX12, whose transcripts enriched in response to *P. vivax* infection(39) in the regulation of changing physiological homeostasis. Sequence conservation of *AsHPX12* with other mosquito species suggested *its* conserved functions in the blood-feeding behavior of hematophagous insects (41). We found *AsHPX12* is constitutively expressed in all the developmental stages of *An. stephensi* mosquitoes designate its possible functions in embryonic cuticle biogenesis and stabilization of extracellular matrix by crosslinking of basal membrane, as reported in *Caenorhabditis elegans* (42).

Previously, a heme-containing salivary secreted peroxidase has been suggested to act as a vasodilator through hydrogen peroxide dependent destruction of serotonin and noradrenalin, but functional role remains uncertain(43). To establish a functional correlation of *AsHPX12* we performed dsRNA mediated silencing and evaluated the altered behavior properties in *An. stephensi* mosquitoes. Surprisingly, an enhanced eagerness of *AsHPX12* knockdown mosquitoes towards a rabbit host, but a significant delay in probing time and probing propensity, together with suggested that HPX12 is key to regulate salivary physiological homeostasis. Additionally, we also noticed that HPX12 silencing causes a significant enrichment of odorant-binding proteins expression in both the salivary gland and olfactory system, but in parallel down-regulate some of the salivary cocktail proteins expression such as Apyrase (also see Supplemental Fig1) which are key to regulate pre-blood meal associated host-seeking properties (44). An earlier study by Sim et al. (2012) showing altered probing time and host-seeking abilities after silencing of salivary OBPs, correlates a possible chemosensory signaling regulation of salivary proteins in the mosquito *Ae. aegypti* (45).

We interpret, cellular disruption and altered expression of salivary proteins caused by HPX12 dysregulation, may have a direct influence on mosquito’s salivary-chemosensory derived host-seeking abilities, affecting the blood meal acquisition process. These observations further corroborate and support the idea that the down-regulation of salivary apyrase is crucial to enhances host attraction of the *Plasmodium*-infected than un-infected mosquitoes (46). Recently, heme-peroxidase homolog HPX15, has been suggested to act as an agonist and likely favor the survival of endogenous gut-bacterial population as well as *Plasmodium* parasite, by the formation of cross-linked mucin barrier on the luminal side of the midgut (40). Elevation of salivary *AsHPX12* mRNA level after 24hrs of blood-feeding and consistent up-regulation in response to salivary invaded *P. vivax* sporozoites, together indicate that *AsHPX12* may have a dual role in the management of salivary gland homeostasis and long term survival of stored sporozoites. Further insights into the sporozoite storage mechanism within the salivary gland may allow us to confirm this hypothesis.

## Conclusion

In summary, for the first, we demonstrate that salivary specific *AsHPX12* plays an important role in the regulation of pre-blood associated behavioral properties. A parallel transcriptional modulation in response to blood feeding and *P. vivax* infection, suggests salivary *AsHPX12* may have a pivotal role in the management of physiological homeostasis, *Plasmodium* sporozoite survival, and transmission.

## Acknowledgment

We would like to thank all the technical staff members of the central insectary for mosquito rearing and Kunwarjeet Singh for lab assistance. We are grateful to the malaria clinic facility support for their contribution *Plasmodium* infection study. Finally, we thank Xceleris Genomics, Ahmedabad for NGS sequencing.

## Authors’ contribution

SK, RKD conceived scientific hypothesis and designed the experiments.CC, JR, ST, TDD, and PS Technical support for tissue dissection/collection and gene profiling, Data reviewing, and presentation.SV provides support for bioinformatics data analysis.RKD, KCP contributed reagents/ materials/ Analysis tools; SK wrote the paper, KCP, RKD edit the MS. All authors read and approved the final manuscript.

