## Supplemental data for "Salivary *AsHPX12* influence pre-blood meal associated behavioral properties in the mosquito *Anopheles stephensi*"

**Table 1**. Percentage identity match of putative *AsHPX12* peroxidases retrieved from different species of Anopheles mosquitoes (major vector, minor vector, and non-vector)

| Vectorial capacity | Species Name | ID | E-value | Score | Identity % |
|---|---|---|---|---|---|
| **Major vectors** | *A. arabiensis* | AARA008970-RA | 0.00 | 545 | 76.6 |
| | *A. atroparvus* | AATE017374-RA | 0.00 | 567 | 67 |
| | *A.culicifacies* | ACUA003065-RA | 0.00 | 543 | 83.5 |
| | *A. darlingi* | ADAC002183-RA | 0.00 | 583 | 55.8 |
| | *A. dirus* | ADIR015871-RA | 0.00 | 1191 | 74.2 |
| | *A. farauti* | AFAF007386-RA | 0.00 | 1038 | 69.68 |
| | *A. gambiae* | AGAP029195-RA | 0.00 | 1110 | 85.7 |
| | *A. maculatus* | AMAM023652-RA | 4e-146 | 436 | 53.8 |
| | *A.sinensis* | ASIS004536-RA | 0.00 | 539 | 74.8 |
| | *A. stephensi* | ASTE016356-RA | 0.00 | 533 | 100 |
| **Minor vectors** | *A.epiroticus* | AEPI000705-RA | 0.00 | 851 | 73 |
| | *A. albimanus* | AALB016040-RA | 0.00 | 1024 | 60.06 |
| | *A.melas* | AMEC003752-RA | 0.00 | 530 | 75.2 |
| | *A.merus* | AXCO02016292 | 0.00 | 528 | 80.2 |
| | *A. minimus* | AMIN015997-RA | 0.00 | 694 | 55.7 |
| **Non-vectors** | *A. christyi* | ACHR005516-RA | 3e-158 | 423 | 52.4 |
| | *A.quadriannulatus* | AQUA002357-RA | 0.00 | 876 | 73.4 |

**Table 2. List of primers used for this study**

| S.No. | Gene Name | Primer Sequence |
|---|---|---|
| 1 | HPX2 | Fw: CACGAAGCTAAAAATTGTCC<br>Rev: AAGAGATGCTCCAGATCGTA |
| 2 | HPX3 | Fw: AGTTCTTCACGGTTCATCAC<br>Rev: GATCTTCTCCAGGTGTTTTG |
| 3 | HPX8 | Fw: AAGCTGTACCAAGAAGCTC<br>Rev: GCTCCTGATACTTCGTTAAA |
| 4 | HPX10 | Fw: AAGAAGGTTGACGGTACAGA<br>Rev: CAACGTTAGCAGTCGTATCA |
| 5 | HPX15 | Fw: CTATTGTCGCAACAGTACGA<br>Rev: CGGTAAACTGTCCATCATTT |
| 6 | Duox | Fw: TACGAGGATTTCAAGCTGAT<br>Rev: GTGTAGCAGTCCCACTTTTC |
| 7 | HPX12 | Fw: GAACAGTGCCACCGATACCT<br>Rev: CCGAGATAATAGGGCAACCA |
| 8 | Apyrase | Fw: CTCAAGAGATTGGGAAGAC<br>Rev: CCTATTTGGATGGGATTAC |
| 9 | D7L | Fw: GATCTATGACCCGCAGGAAA<br>Rev: ATCGGTTGTGAACTGGAAGG |
| 10 | 53.7kDa | Fw: TACAGCTAGCGGCAAGGAAT<br>Rev: AGGTAACGGACAGTGCCATC |
| 11 | DsrLacz | Fw: 5' TAATACGACTCACTATAGGGGAGTCAGTGAGCGAGGAAG 3'<br>Rev: 5'TAATACGACTCACTATAGGGTATCCGCTCACAATTCCACA 3' |
| 12 | DsrHPX12 | Fw: 5' TAATACGACTCACTATAGGGTTCTGGTGTTTGCCATCGTA 3'<br>Rev: 5' TAATACGACTCACTATAGGGCAGGATGTTCTGCTCGTTGA 3' |

| | | |
|---|---|---|
| 13 | Anopheline | Fw: GCGAGGAGCCTGAATATGAC<br>Rev: AATCCGACTGGTTTTCGTTG |
| 14 | 37.3kDa | Fw: ACACACGCAAAATGTGCTTC<br>Rev: ATCAAAGTCTCCACCGTTCG |
| 15 | SG2B | Fw: GTCGGGTGGATTTGGATTG<br>Rev: TTAACCGAAAAATGGGAAACC |
| 16 | Hypothetical 16 | Fw: TGTGTGATCGTCAAAAACTC<br>Rev: TTGCCATTCTTATCGATTTC |
| 17 | OBP10 | Fw: AAGTGAAGGGATACAAGCTG<br>Rev: CGTATGTCACACTGCTTCAC |
| 18 | OBP20 | Fw: GTTACGTGAACTGCGTGAT<br>Rev: GTTCTTCGAAAGGCACTGA |
| 19 | Actin | Fw: TGCGTGACATCAAGGAGAAG<br>Rev: GATTCCATACCCAGGAACGA |
| 21 | IR75K | Fw: GGAAGATACGGTTTACAATC<br>Rev: GATGACCTTCAGATATTCCA |

**Table 3:** Table showing comparison of host seeking behavior for blood feeding of naïve mosquito and HPX12 silenced mosquito in three biological replicates.

| SET | Date | Control | HPX-12 dsr injected | Total mosquito |
|---|---|---|---|---|
| 1. | 29/5/18 | 7 | 20 | 25 |
| 2. | 9/7/2018 | 17 | 22 | 30 |
| 3. | 18/7/2018 | 28 | 46 | 60 |
| | Total | 52 | 88 | 115 |
| | Percent | 45% | 76% | |

**Table 5**: String interaction in tabular form: containing node annotation and score

| Node1 | Node2 | Node1 accession | Node2 accession | Node1annotation | Node2 annotation | score |
|---|---|---|---|---|---|---|
| 11175919 | 1276489 | AGAP013192-PA | AGAP005822-PA | *Venom allergen* | *annotation not available* | 0.159 |
| 11175919 | 1277100 | AGAP013192-PA | AGAP006506-PA | *Venom allergen* | *annotation not available* | 0.152 |
| 11175919 | 1281804 | AGAP013192-PA | AGAP001374-PA | *Venom allergen* | *annotation not available* | 0.151 |
| 11175919 | AGAP009974-PA | AGAP013192-PA | AGAP009974-PA | *Venom allergen* | *annotation not available* | 0.153 |
| 11175919 | AGAP010735-PA | AGAP013192-PA | AGAP010735-PA | *Venom allergen* | *annotation not available* | 0.265 |
| 1270958 | 1274294 | AGAP011026-PA | AGAP003629-PA | *5' nucleotidase, ecto; Belongs to the 5'-nucleotidase family* | *AGAP003629-PA; 5'-nucleotidase ; Belongs to the 5'-nucleotidase family* | 0.170 |
| 1270958 | 1276489 | AGAP011026-PA | AGAP005822-PA | *5' nucleotidase, ecto; Belongs to the 5'-nucleotidase family* | *annotation not available* | 0.354 |
| 1270958 | 1277100 | AGAP011026-PA | AGAP006506-PA | *5' nucleotidase, ecto; Belongs to the 5'-nucleotidase family* | *annotation not available* | 0.349 |
| 1270958 | 1277947 | AGAP011026-PA | AGAP008004-PA | *5' nucleotidase, ecto; Belongs to the 5'-nucleotidase family* | *annotation not available* | 0.288 |
| 1270958 | 1281804 | AGAP011026-PA | AGAP001374-PA | *5' nucleotidase, ecto; Belongs to the 5'-nucleotidase family* | *annotation not available* | 0.349 |
| 1270958 | 4577351 | AGAP011026-PA | AGAP001192-PA | *5' nucleotidase, ecto; Belongs to the 5'-nucleotidase family* | *annotation not available* | 0.280 |
| 1270958 | AGAP009974-PA | AGAP011026-PA | AGAP009974-PA | *5' nucleotidase, ecto; Belongs to the 5'-nucleotidase family* | *annotation not available* | 0.532 |
| 1270958 | AGAP010735-PA | AGAP011026-PA | AGAP010735-PA | *5' nucleotidase, ecto; Belongs to the 5'-* | *annotation not available* | 0.362 |

| | | | | | | |
|---|---|---|---|---|---|---|
| | | | | *nucleotidase family* | | |
| 1274294 | 1270958 | AGAP003629-PA | AGAP011026-PA | *AGAP003629-PA; 5’-nucleotidase ; Belongs to the 5’-nucleotidase family* | *5’ nucleotidase, ecto; Belongs to the 5’-nucleotidase family* | 0.170 |
| 1274294 | 1276489 | AGAP003629-PA | AGAP005822-PA | *AGAP003629-PA; 5’-nucleotidase ; Belongs to the 5’-nucleotidase family* | *annotation not available* | 0.288 |
| 1274294 | 1277100 | AGAP003629-PA | AGAP006506-PA | *AGAP003629-PA; 5’-nucleotidase ; Belongs to the 5’-nucleotidase family* | *annotation not available* | 0.288 |
| 1274294 | 1277947 | AGAP003629-PA | AGAP008004-PA | *AGAP003629-PA; 5’-nucleotidase ; Belongs to the 5’-nucleotidase family* | *annotation not available* | 0.349 |
| 1274294 | 1281804 | AGAP003629-PA | AGAP001374-PA | *AGAP003629-PA; 5’-nucleotidase ; Belongs to the 5’-nucleotidase family* | *annotation not available* | 0.286 |
| 1274294 | 4577351 | AGAP003629-PA | AGAP001192-PA | *AGAP003629-PA; 5’-nucleotidase ; Belongs to the 5’-nucleotidase family* | *annotation not available* | 0.286 |
| 1274294 | AGAP009974-PA | AGAP003629-PA | AGAP009974-PA | *AGAP003629-PA; 5’-nucleotidase ; Belongs to the 5’-nucleotidase family* | *annotation not available* | 0.304 |

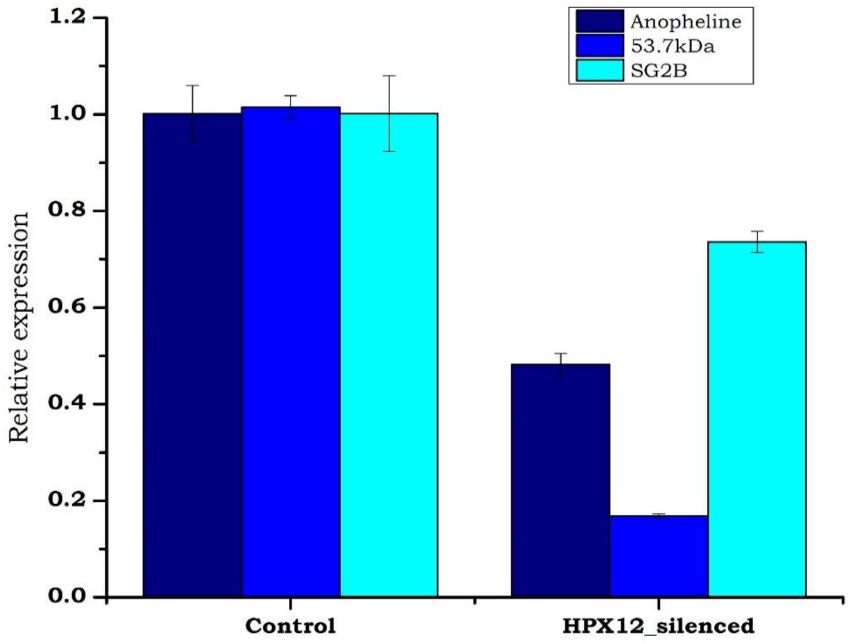


**Figure S1:** Alteration of salivary gland cocktail proteins in *AsHPX12* knock down mosquitoes. Data from three independent experiments using separate generations of mosquitoes were pooled.
